# Characterization of Single Protein Dynamics in Cell Plasma Membrane Derived Polymer Cushioned Lipid Bilayers

**DOI:** 10.1101/641258

**Authors:** Wai Cheng (Christine) Wong, Jz-Yuan Juo, Yi-Hung Liao, Ching-Ya Cheng, Chih-Hsiang Lin, Chia-Lung Hsieh

## Abstract

Native cell membrane derived supported lipid bilayers (SLBs) are emerging platforms that have broad applications ranging from fundamental research to next-generation biosensors. Central to the success of the platform is proper accommodation of membrane proteins so that their dynamics and functions are preserved. Polymer cushions have been commonly employed to avoid direct contact of the bilayer membrane to the supporting substrate, and thus the mobility of transmembrane proteins is maintained. However, little is known about how the polymer cushion affects the absolute mobility of membrane molecules. Here, we characterized the dynamics of single membrane proteins in polymer-cushioned lipid bilayers derived from cell plasma membranes and investigated the effects of polymer length. Three membrane proteins of distinct structures, i.e., GPI-anchored protein, single-pass transmembrane protein CD98 heavy chain, and seven-pass transmembrane protein SSTR3, were fused with green fluorescence proteins (GFPs) and their dynamics were measured by fluorescence single-molecule tracking. An automated data acquisition was implemented to study the effects of PEG polymer length to protein dynamics with large statistics. Our data showed that increasing the PEG polymer length (molecular weight from 1,000 to 5,000) enhanced the mobile fraction of the membrane proteins. Moreover, the diffusion coefficients of transmembrane proteins were raised by increasing the polymer length, whereas the diffusion coefficient of GPI-anchored protein remained almost identical with different polymer lengths. Importantly, the diffusion coefficients of the three membrane proteins became identical (2.5 μm^2^/s approximately) in the cushioned membrane with the longest polymer length (molecular weight of 5,000), indicating that the SLBs were fully suspended from the substrate by the polymer cushion at the microscopic length scale. Transient confinements were observed from all three proteins, and increasing the polymer length reduced the tendency of transient confinements. The measured dynamics of membrane proteins were found to be nearly unchanged after depletion of cholesterol, suggesting that the observed immobilization and transient confinement were not due to cholesterol-enriched membrane nanodomains (lipid rafts). Our single-molecule dynamics elucidate the biophysical properties of polymer cushioned plasma membrane bilayers that are potentially useful for future developments of membrane-based biosensors and analytical assays.

## INTRODUCTION

Biological membranes are heterogeneous both in composition and in spatial organization. Plasma membranes of mammalian cells are comprised of hundreds of different lipids and proteins.^1-3^ The mixture of diverse molecular species that differ in their physical and chemical properties is expected to be heterogeneous on the molecular length scale (1–10 nm).^4^ Meanwhile, most membrane functions start from the interaction between molecules at the microscopic length scale. Therefore, mapping the membrane organization at the molecular scale is useful for understanding the local environment and condition for membrane activity.^5^ Furthermore, biological membranes are fluidic where most molecules move constantly in the membrane driven by thermal fluctuation. The membrane fluidity facilitates stochastic molecular interplays that are essential for many membrane functions.^6-7^

Optical microscopy is arguably the most powerful tool for studying membrane organization and dynamics in living organisms.^8-10^ With proper labeling, single membrane molecules, such as lipids and proteins, can be visualized and tracked at high resolutions in live cells.^8-11^ A lot has been learned from single-molecule tracking (SMT) in the plasma membranes. For example, nanoscopic membrane compartmentalization,^11^ transient trapping of lipids^12^ and protein-protein interactions^13^ were revealed in live cells by high-speed SMT. The unique advantage of SMT is to measure dynamics of individual molecules without ensemble average, allowing for detecting heterogeneous behaviors in a large molecular population.^8^ Although it is promising to study membrane dynamics in live cells, the measurements and data interpretation have often been complicated by the complexity of the cells, making it difficult to probe the intrinsic molecular behaviors of the membrane. For example, in the plasma membranes of live cells, molecular dynamics are critically affected by the actin filaments underneath the membrane whose effect is difficult to estimate quantitatively.^14-15^ Without accounting the effect of actin filaments properly, quantitative measure of inherent membrane dynamics and molecular interactions becomes unattainable. Another difficulty in live cell experiments is the inevitable heterogeneity in cell states and cell types, which may bias the results of membrane dynamics.^16-17^ Moreover, cells undergo continuous endocytic cycles through endocytosis and exocytosis, leading to active transport of membrane that further complicates the study of molecular dynamics in membranes.^18^ Finally, cells produce optical background signal (e.g., autofluorescence) that makes single-molecule detection and imaging more difficult.

In many cases, it is easier or even more informative to study intrinsic membrane properties (dynamics and organization) in an in vitro system without living cells.^19-28^ Several cell-free membrane platforms have been demonstrated with different degrees of complexity that can mimic the cell membranes to various levels. For example, membrane proteins were reconstituted into model membranes where their dynamics and functions were examined.^29-34^ Such membrane protein reconstitution was limited to a rather simple molecular composition that is comprised of a few different membrane protein and lipid species. To mimic the complex composition of native cell membranes, giant plasma membrane vesicle (GPMV), a cell-derived vesicle of a diameter of tens of micrometers, is a widely used as a cell-free system for studying various membrane properties, including membrane dynamics, phase separation, and protein behaviors.^24-25, 35^ The GPMV largely preserves the membrane composition and molecular orientation, making it a unique platform to study membrane proteins in native environments. Unfortunately, the spherical shape of the GPMV is less compatible with optical microscope observation and other surface-based analytical methods.

Recently, there have been a few demonstrations of preparing planar supported lipid bilayers (SLBs) by native cell membranes.^36-40^ The SLBs are planar membrane platform that is highly stable and suitable for optical microscope measurements.^22, 41-43^ There are several key advantages of studying biomembranes in SLBs derived from native cell membranes. First, the membrane proteins readily reside in the bilayer membranes without the need of detergent-based extraction, purification and reconstitution, which greatly minimizes the possible perturbation to protein structure and function.^44^ Second, the SLBs derived from native cell membranes potentially support supramolecular complexes that are closer to their original configuration in the cell membranes, giving a better chance to reproduce molecular interactions in the membrane of live cells. Preparing SLBs derived from native cell membranes, however, is not trivial. The SLBs are typically prepared from liposomes that are mostly comprised of synthetic lipids, and the formation of SLBs relies on self-rupture and fusion of liposomes on a hydrophilic supporting surface.^45^ On the other hand, native cell plasma membrane vesicles normally do not rupture when adsorbing on a substrate. To resolve this issue, it has been demonstrated that one can trigger the rupture and fusion of cell plasma membrane vesicles by adding high-concentration of polymer solution^46^ or synthetic liposomes^36^ during the SLB preparation. Moreover, by enhancing the vesicle-substrate electrostatic interaction, supported cell plasma membranes were also successfully prepared.^38^ Importantly, by following a protocol described by Daniel group, it was shown that the orientation of membrane proteins can be maintained in the SLB platform.^37^

In addition to the membrane formation, a central concern about the supported plasma membranes is whether the mobility and function of membrane protein are preserved. Conventional SLBs have a thin water layer (1–2 nm) between the membrane and the supporting substrate,^47^ which is too narrow for accommodating the cytoplasmic domains of most transmembrane proteins. The contact of transmembrane protein with the substrate not only immobilizes but also potentially denatures the protein.^48^ Several methods have been demonstrated to create a space between the membrane and the substrate.^22^ Among these methods, polymer cushion appears to be a reliable approach which has successfully produced SLBs with mobile and functional reconstituted membrane proteins.^22, 30, 32-34, 48-51^ Although it is encouraging to see the protein mobility and some capability of ligand binding are preserved in polymer-cushioned SLBs,^30, 32-33^ it remains elusive how the polymer cushion determines the absolute mobility of membrane protein. We point out that the diffusion coefficients of membrane molecules (e.g., lipids and GPI-anchored fluorescent proteins) measured in polymer-cushioned SLBs derived from native cell membranes are slower than those measured in GPMVs,^37-38, 52^ indicating the membrane fluidity is affected by the cushioned substrate. Finally, larger proteins seem to diffuse more slowly, but the underlying mechanism is still unclear.^37-38^

In this work, we study the effect of polymer cushion to diffusion characteristics of membrane proteins in SLBs prepared with cell plasma membrane vesicles. Three different polymer lengths, giving three different spacing between the SLB and the substrate, are investigated. For each polymer length, we characterize diffusion of three types of membrane proteins fused with green fluorescent protein (GFP) by fluorescence single-molecule imaging and tracking. The three membrane proteins are peripheral protein (GPI-anchored GFP),^53^ single-pass transmembrane glycoprotein (heavy subunit protein of CD98),^54^ and seven-pass G-protein coupled receptor SSTR3 (Shekel Somatostatin receptor type 3).^55^ The three proteins are different in size and also in their affinities to cholesterol-dependent lipid rafts.^56^ We further evaluate the effect of lipid rafts to their diffusion by modulating the cholesterol concentration in the membrane.

## MATERIALS AND METHODS

### Materials

1-palmitoyl-2-oleoyl-glycero-3-phosphocholine (16:0-18:1 POPC), 1,2-dipalmitoyl-sn-glycero-3-phosphoethanolamine-N-[methoxy(polyethylene glycol)-1000] (16:0 PEG1000-DPPE), 1,2-dipalmitoyl-sn-glycero-3-phosphoethanolamine-N-[methoxy(polyethylene glycol)-3000] (16:0 PEG3000-DPPE) and 1,2-dipalmitoyl-sn-glycero-3-phosphoethanolamine-N-[methoxy(polyethylene glycol)-5000] (16:0 PEG5000-DPPE) were purchased from Avanti Polar Lipids, Inc. Atto532-DOPE was purchased from ATTO-TEC GmbH. The GPI-GFP plasmid was a generous gift from Ilya Levental of the University of Texas. The plasmid of GFP-fused CD98 heavy chain was purchased from GeneCopoeia™ (EX-G00009-M98). The plasmid of GFP-fused SSTR3 was obtained from Addgene (pEF5B-FRT-AP-Sstr3-GFP-DEST, a gift from Maxence Nachury; Addgene plasmid # 49098).^57^

### Cell culture and transfection

HeLa cells were maintained in Minimum Essential Medium (Hyclone™) supplemented with 10% fetal bovine serum (Hyclone™) and 100 U/mL penicillin and 10 µg/mL streptomycin (Hyclone™). 8 × 10^5^ HeLa cells per dish were seeded in 10 cm culture dishes (Corning) pre-coated with Poly-D-Lysine (Sigma-Aldrich, Merck) and incubated for 24 h at 37 °C supplemented with 5% CO_2_. 30 µL of PEI transfection reagent (Sigma-Aldrich, Merck) and 10 µg of DNA plasmid, pGFP-GPI, pEF5B-FRT-AP-Sstr3-GFP-DEST or CD98-GFP, were used for transfection for each dish. Successful transfection was confirmed by fluorescence confocal microscopy (Figure S1). Cells were grown for 24 h at 37 °C incubator supplemented with 5% CO_2_.

### Preparation of plasma membrane vesicles

Plasma membrane vesicles were induced chemically based on an established protocol.^35^ Briefly, growth medium was removed and each dish was rinsed with PBS, followed by GPMV buffer (2 mM CaCl_2_, 10 mM HEPES, 150 mM NaCl at pH 7.4). Then, 4 mL of GPMV buffer with 25 mM paraformaldehyde (PFA) and 2 mM dithiothreitol (DTT) was added to each dish. Cells were incubated at 37 °C, 5% CO_2_ for 1-2 h. Plasma membrane vesicles were collected and cell debris was removed with 0.22 µm syringe filter unit (Millex-GV Syringe Filter Unit, 0.22 µm, PVDF, 33 mm, Merck). Vesicles were stored in aliquots at 4 °C until use.

### Preparation of PEGylated liposomes

Four types of lipid mixture were used in this work, namely POPC (100% POPC), PEG1000 (99.5% POPC 0.5% PEG1000-DPPE), PEG3000 (99.5% POPC 0.5% PEG3000-DPPE) and PEG5000 (99.5% POPC 0.5% PEG3000-DPPE). Lipids were mixed in chloroform at desired molar ratio. Chloroform was removed by gentle blowing with nitrogen gas followed by desiccation in vacuum for at least 1 h to remove residual chloroform. Dried lipid films were hydrated in PBS (150 mM NaCl, 5 mM NaH_2_PO_4,_ 5 mM Na_2_HPO_4_ at pH 7.4) at 1 mg/mL for 1 hr at room temperature. Lipid solution was then vortexed and 4-fold diluted before sonication using a pre-cleaned tip sonicator at 4 °C (Q700, Qsonica, CT, USA). Suspension was carefully transferred to Eppendorf tubes, 1 mL each and centrifuged at 16,000 × g for 20 min to precipitate lipid debris. Supernatant was carefully transferred to new Eppendorf tubes and stored at 4 °C before use.

### Formation of SLBs with plasma membrane vesicles and synthetic liposomes

SLBs were formed using a method described previously with slight modification.^37^ Briefly, 8-well chambered coverglass (Nunc Lab-Tek II, Thermo Scientific, MT, USA) were cleaned by incubating in 2% Hellmanex, 1M KOH, and deionized water sequentially for 15 mins each. Then, 150 µL of 2.8 × 10^8^ plasma membrane vesicles/mL were deposited into each well and incubated for 15 min to ensure sufficient amount of vesicles were adsorbed onto glass surface. Wells were rinsed with PBS (5 mM NaH_2_PO_4_, 5 mM Na_2_HPO_4_, 150 mM NaCl at pH 7.4) to remove unadsorbed vesicles. 150 µL of 7 × 10^8^ liposomes/mL were added and incubated for at least 30 min to allow fusion of liposomes with plasma membrane vesicles. Wells were then rinsed thoroughly with PBS to remove excess liposomes.

### Cholesterol depletion

Cholesterol was depleted using an established protocol with slight modification.^58^ Briefly, desired amount of methyl-β-cyclodextrin (MβCD) powder is dissolved in deionized water to a final concentration of 3 mM and supplemented with 25 mM HEPES. Sample was incubated with MβCD solution for at least 15 min at room temperature prior to imaging.

### Single-molecule fluorescence microscope imaging

For imaging and tracking single GFP-fused membrane proteins, we used an inverted microscope (IX83, OLYMPUS) with addition of a laser (OBIS 488 nm, Coherent) for illumination. Excitation intensity was kept at 2–3 kW/cm^2^ during image acquisition. Fluorescence signal was collected by an oil-immersion microscope objective (UPLSAPO 100X, NA 1.4 Olympus) and imaged on to an EMCCD (iXon Ultra 897, Andor). Videos were recorded at 125 Hz (6.24 ms exposure time) with an image resolution of 146 × 146 pixels and pixel size of 158 nm. An automated data acquisition procedure was set up by synchronizing the laser illumination, lateral translation of the sample, and video recording. First, the sample was brought to the focal plane of the objective by auto-focusing without laser excitation. A continuous video recording of 250 frames was initiated and synchronized with laser excitation. After recording the 250-frame video, sample was laterally translated to a fresh area without laser illumination for the next cycle of video recording. Our procedure allows for non-biased data acquisition that captures both the mobile and immobile membrane molecules. 600 videos were recorded for each sample, covering a sample area over 500 × 500 μm^2^. Fluorescence imaging of ATTO532-DOPE was performed in a similar setup but with a 532 nm laser light source.

### Single-particle tracking

A Gaussian smoothing was performed on every fluorescence image to reduce electronic noise. A bright spot in the smoothened image was considered as a protein when its intensity is above certain threshold. The signal-to-noise ratio (SNR) of a single GFP is approximately 5, which is sufficient for reliable detection. To determine the position of the protein with sub-pixel accuracy, a 5 × 5 sub-image containing the bright spot was cropped and fitted by a 2D Gaussian function with five fitting parameters: amplitude, two lateral widths and positions. The localization precision can be estimated directly from the quality of fitting,^59^ which is typically 20 nm in our measurements. A trajectory is obtained by connecting the nearest neighboring bright spots in the consecutive frames.

### Estimation of diffusion coefficient

We used trajectories longer than 25 steps for estimation of diffusion coefficient. The mean square displacement (MSD) of individual trajectory was calculated. The diffusion coefficient is best estimated by the slope of the first two MSD data points because our localization precision and frame time are sufficiently high.^60^

### Detection of transient confinement

To quantify the protein confinement behavior, we followed the method developed previously.^61^ We first calculated the probability level (L) of segments cropped from each particle trajectory by using equations: 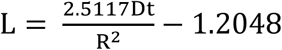, where t is the time lapse of the segment; R is the radius for a segment determined by the point in the segment with the largest displacement from the starting point; D is the diffusion coefficient estimated from all recorded trajectories of the mobile fraction of a given protein. A segment of trajectory with L above a certain threshold (Lc) was characterized as a confinement zone. Specifically, we chose the values of segment size (Sm=13) and threshold (Lc=55) which enables reliable detection of confined motion in the trajectory. Using the same detection criteria, we found nearly no confinement (around 0.2%) in simulated Brownian trajectories with the same number of steps and localization error of experimental data. We confirmed that our results remain largely the same when fine-tuning the values of Sm and Lc, indicating the conclusions were not sensitive to these parameters.

## RESULTS

### Single molecule diffusion in polymer cushioned plasma membrane bilayers

Our supported bilayer membrane was prepared by mixing cell plasma membrane vesicles and synthetic liposomes on clean coverglass (see detailed protocol in Methods). The size of cell plasma membrane vesicles and synthetic liposomes were both around 150 nm, characterized by nanoparticle tracking analysis (NTA, Nanosight NS300, Malvern, Figure S2). A small amount (0.5 mol%) of PEGylated lipid was added in the synthetic liposome that forms a polymer cushion spontaneously after rupture and fusion.^37^ The density of PEGylated lipid is low and it is expected to appear in the form of “mushroom” that levitates the membrane,^48^ creating a space (a few nanometers depending on the PEG length) between the membrane and the substrate for accommodating the transmembrane proteins (Figure 1a). For optical observation, GFP was fused to membrane protein of interest that was introduced to the membrane through plasma membrane vesicle by transfection (Methods). Single-molecule epi-fluorescence microscope imaging was performed to visualized dynamics of individual GFP-fused proteins (Methods). Three membrane proteins, i.e., GPI-GFP, GFP-fused CD98 heavy chain (CD98-GFP), and GFP-fused SSTR3 (SSTR3-GFP), were studied and their schematics are plotted in Figure 1a.

**Figure 1.**
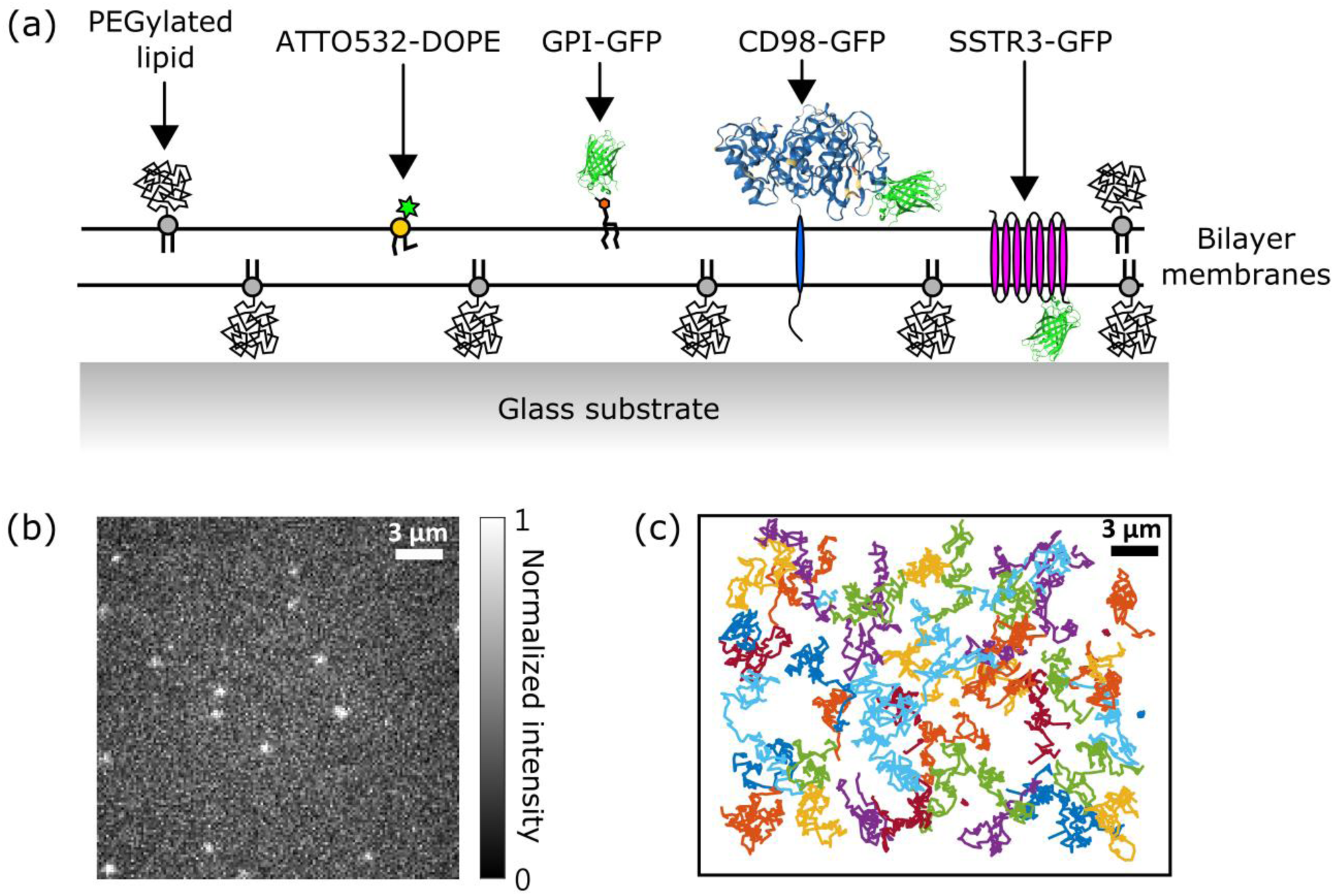
Single-molecule fluorescence imaging and tracking in polymer cushioned plasma membrane bilayers. (a) Schematics of the membrane molecules studied in this work. The ATTO532-DOPE is a dye-labeled phospholipid. The GPI-GFP is a glycolipid, glycophosphatidylinositol (GPI), with a GFP attached in the headgroup. The CD98-GFP heavy chain is a single-pass transmembrane protein and is fused with GFP in its C-terminus in extracellular domain. The SSTR3-GFP is a seven-pass transmembrane protein and is fused with GFP in its C-terminus in cytoplasmic domain. (b) A snapshot of fluorescence image of single GPI-GFPs. (c) Representative trajectories of GPI-GFPs in polymer cushioned plasma membrane bilayers.

The GFP-labeled proteins appeared as bright spots in the fluorescence image (Figure 1b). Individual bright spots were confirmed to be single GFPs by their single-step photobleaching (see Movie S1–S3 for GPI-GFP, CD98-GFP, and SSTR3-GFP, respectively). Trajectories were obtained from the fluorescence videos by single-particle tracking (Figure 1c and Methods). From every trajectory that is longer than 25 steps, a diffusion coefficient is estimated by calculating the MSD (Methods). In all membrane samples measured in our study, we observed two distinct diffusion characteristics. There appeared to be a fast diffusing population (diffusion coefficient 1–5 μm^2^/s) accompanied by a slowly diffusing one (diffusion coefficient on the order of 0.01 μm^2^/s). The two populations can be well described by superposition of two Gaussian functions in the histogram plotted in logarithmic scale (data and quantitative analyses are presented in the following sections). The slow diffusion coefficient of 0.01 μm^2^/s matches well with the value of σ2/Δt where σ is the localization error and Δt is the frame time of our fluorescence measurements, indicating the slow diffusion corresponds to immobilization. Thus, we consider the slowly diffusing proteins to be immobile in the rest of the text. Here we emphasize that an automated and high-throughput imaging procedure was implemented (Methods) in order to measure the true dynamics of all proteins without bias (especially caused by photobleaching). Using our methods, at least 1,000 trajectories (longer than 25 steps) were recorded for each membrane protein sample, providing large statistics for investigation of protein dynamics.

### Effect of polymer length

We characterized membrane protein diffusion in three polymer cushioned plasma membrane bilayers with different PEG lengths, prepared by adding PEG1000-DPPE, PEG3000-DPPE, and PEG5000-DPPE in the bilayers, respectively (Methods). We first probed the membrane fluidity by measuring the diffusion of a dye-labeled lipid, ATTO532-DOPE, that was introduced to the membrane through synthetic liposomes. Movie S4 shows the fluorescence video of ATTO532-DOPE in a PEG1000 membrane. The histograms of diffusion coefficient of ATTO532-DOPE measured in the three membrane samples are plotted in Figure 2a. Most lipids diffused fast with a diffusion coefficient of around 3 μm^2^/s, and only a small fraction (no more than 16%) of lipids appears to be immobile. Increasing the PEG length slightly reduces the immobile fraction. In the membrane of PEG5000, the immobile fraction is as low as 7 %. Interestingly, the diffusion coefficient of lipid is nearly independent of the polymer length. We measured 3.0 ± 0.9, 2.9 ± 0.9, and 2.9 ± 0.8 μm^2^/s (mean ± standard deviation) in PEG1000, PEG3000, and PEG5000 membranes, respectively.

**Figure 2.**
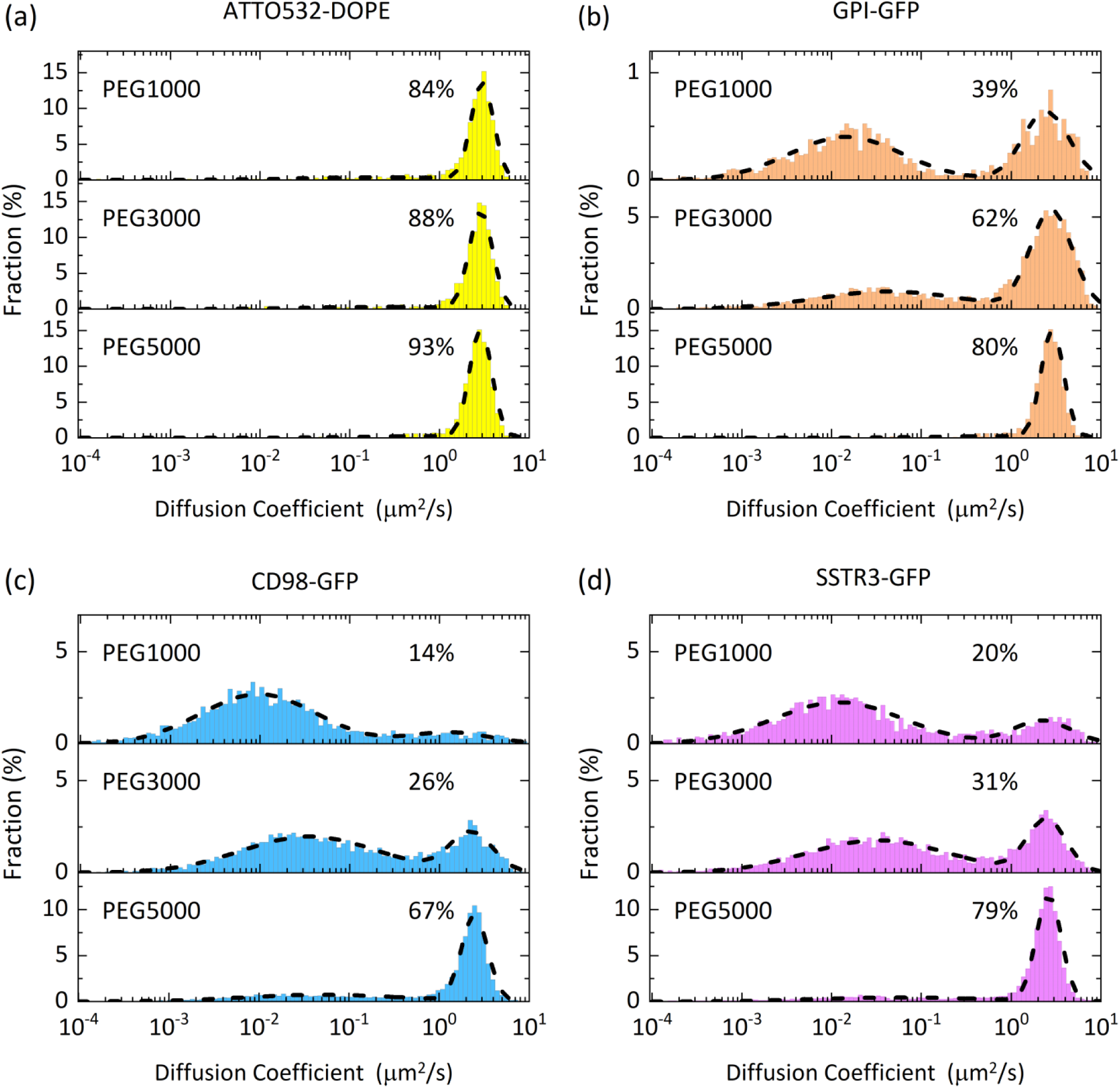
Histograms of diffusion coefficient of four membrane molecules (a) ATTO532-DOPE, (b) GPI-GFP, (c) CD98-GFP, and (d) SSTR3-GFP in polymer cushioned plasma membrane bilayers with different polymer lengths. Histograms were fitted by a superposition of two Gaussian functions, plotted in dashed lines. The percentage displayed in each histogram marks the mobile fraction of the molecule.

Same measurements were performed on GPI-GFP that is a lipid-anchored protein. Histograms of diffusion coefficient of GPI-GFP were plotted in Figure 2b. In the PEG1000 membrane, significant amount (61 %) of GPI-GFP was immobile. Movie S1 shows the fluorescence video of GPI-GFP in a PEG1000 membrane. We note that our membrane preparation is expected to preserve the membrane orientation,^37^ and the GPI-GFP should reside in the outer leaflet of the bilayer, facing the bulk aqueous solution (see Figure 1a). Without direct contact to the substrate, the GPI-GFP is expected to be immobilized through interleaflet coupling and molecular pinning.^62^ The other 39 % of GPI-GFP diffused fast with a diffusion coefficient of 2.4 ± 1.5 μm^2^/s. By increasing the PEG length, the mobile fraction of GPI-GFP was enhanced—62 % in PEG3000 membrane and 80 % in PEG5000 membrane. The enhancement of the mobile fraction suggests that PEG cushion effectively reduces the strength of interleaflet coupling and thus the molecular pinning. For the mobile population, we measured similar diffusion coefficients of GPI-GFP in the three membranes, i.e., 2.4 ± 1.5, 2.8 ± 1.5, and 2.5 ± 0.9 μm^2^/s (mean ± std) in PEG1000, PEG3000, and PEG5000 membranes, respectively. The diffusion of GPI-GFP is slower than that of ATTO532-DOPE. The slower diffusion of GPI-GFP may be due to its larger headgroup and its distinct affinity to cholesterol dependent lipid nanodomains (lipid rafts). The effect of cholesterol will be examined in the later section.

We then measured the diffusion of two transmembrane proteins, CD98-GFP and SSTR3-GFP. The histograms of diffusion coefficient of CD98-GFP and SSTR3-GFP were plotted in Figure 2c and 2d, respectively. The two proteins appeared to be largely immobile in PEG1000 membrane (86 % for CD98 and 80% for SSTR3). Increasing the PEG length rescued the mobility of the two proteins. The mobile fractions of CD98-GFP (SSTR3-GFP) were 14 (20), 26 (31), and 67 (79) % in PEG1000, PEG3000, and PEG5000 membranes, respectively. Importantly, we found that the diffusion coefficient for the mobile population of CD98-GFP and SSTR3-GFP was enhanced when the PEG length increased: the diffusion coefficients of the mobile CD98-GFP (SSTR3-GFP) were measured to be 1.3 ± 0.5 (2.2 ± 1.7), 2.3 ± 1.3 (2.5 ± 1.3), and 2.5 ± 0.8 (2.6 ± 1.9) μm^2^/s in PEG1000, PEG3000, and PEG5000 membranes, respectively. Such dependency of diffusion coefficient on PEG length was not observed in ATTO532-DOPE and GPI-GFP. We note that CD98-GFP and SSTR3-GFP are transmembrane proteins, while ATTO532-DOPE and GPI-GFP are lipid and lipid-anchored peripheral protein. Our data imply that the diffusion coefficient of transmembrane protein is sensitive to the cushion PEG length possibly because the introduction of PEG cushion helps to reduce the nonspecific interaction of the cytoplasmic domain of transmembrane protein with the supporting substrate. On the other hand, GPI-GFP resides in the distal membrane leaflet where the diffusion coefficient is not sensitive to the PEG cushion. We further point out that although CD98 heavy chain is a single-pass membrane protein, it is expected to form heterodimer with CD98 light chain which is a twelve-pass membrane protein.^63^ As a result, the measured CD98-GFP dynamics is likely to reflect the motion of CD98 heterodimer. The CD98 heterodimer has larger transmembrane domains than SSTR3, which may explain the slower diffusion measured from CD98-GFP than SSTR3-GFP. Table 1 summarizes the diffusion characteristics of all membrane molecules studied in all samples of this work.

**Table 1.** Summary of diffusion characteristics of four membrane molecules measured in three polymer cushioned supported plasma membrane bilayers before and after cholesterol depletion.

|  |  | ATTO532-DOPE |  | GPI-GFP |  | CD98-GFP |  | SSTR3-GFP |  |
| --- | --- | --- | --- | --- | --- | --- | --- | --- | --- |
|  |  | before | after | before | after | before | after | before | after |
|  |  | MβCD | MβCD | MβCD | MβCD | MβCD | MβCD | MβCD | MβCD |
| PEG1000 | Mobile Fraction (%) | 84 | 92 | 39 | 34 | 14 | 12 | 20 | 13 |
| | Diffusion coefficient<sup>†</sup><br>($\mu\text{m}^2/\text{s}$) | $3.0 \pm 0.9$ | $3.2 \pm 0.9$ | $2.4 \pm 1.5$ | $2.6 \pm 1.6$ | $1.3 \pm 0.5$ | $1.3 \pm 1.3$ | $2.2 \pm 1.7$ | $1.8 \pm 1.5$ |
|  | Transient confinement<sup>‡</sup><br>(%) | 4.8 | 4.2 | 5.1 | 6.6 | 16.2 | 11.9 | 12.3 | 12.4 |
|  | Number of trajectories | 646 | 897 | 1983 | 1003 | 2927 | 2646 | 2444 | 1120 |
| PEG3000 | Mobile Fraction (%) | 88 | 91 | 62 | 59 | 26 | 18 | 31 | 30 |
| | Diffusion coefficient<sup>†</sup><br>($\mu\text{m}^2/\text{s}$) | $2.9 \pm 0.9$ | $3.1 \pm 0.9$ | $2.8 \pm 1.5$ | $2.8 \pm 1.4$ | $2.3 \pm 1.3$ | $2.4 \pm 1.1$ | $2.5 \pm 1.3$ | $2.9 \pm 1.5$ |
|  | Transient confinement<sup>‡</sup><br>(%) | 4.6 | 1.9 | 4.7 | 4.9 | 12.6 | 23.2 | 9.5 | 11 |
|  | Number of trajectories | 776 | 1243 | 6087 | 1832 | 6857 | 4298 | 4747 | 1262 |
| PEG5000 | Mobile Fraction (%) | 93 | 94 | 80 | 88 | 67 | 78 | 79 | 79 |
| | Diffusion coefficient<sup>†</sup><br>($\mu\text{m}^2/\text{s}$) | $2.9 \pm 0.8$ | $2.9 \pm 0.9$ | $2.5 \pm 0.9$ | $2.6 \pm 1.0$ | $2.5 \pm 0.8$ | $2.9 \pm 0.9$ | $2.6 \pm 1.9$ | $2.6 \pm 0.9$ |
|  | Transient confinement<sup>‡</sup><br>(%) | 0.8 | 0.9 | 3.3 | 2.2 | 5.1 | 3.8 | 3.0 | 3.2 |
|  | Number of trajectories | 1278 | 1457 | 10518 | 8181 | 12167 | 9422 | 8931 | 9189 |

<sup>†</sup> Diffusion coefficient of the mobile population, mean ± standard deviation, determined by Gaussian fitting in the histogram.
<sup>‡</sup> Percentage of transient confinement is defined as the ratio of total residence time of all detected transient confinements to overall observation time of all mobile molecules.

It was unexpected to see similar diffusion coefficients (around 2.5 μm^2^/s) of GPI-GFP, CD98-GFP, and SSTR3-GFP measured in PEG5000 cushioned plasma membranes because these proteins are different in many aspects, including their physical sizes. The size-insensitive diffusion coefficients suggest that these proteins resided in suspended bilayers supported by mushroom-like pillars of PEG polymer where the diffusion coefficient of a membrane inclusion is weakly dependent on its size (in logarithmic dependency), based on Saffman–Delbrück model.^64^ To estimate the size of PEG cushion, it is useful to calculate its Flory radius *R*_F_ in a simplified situation of ideal solvent.^48, 65-66^ Given the PEG subunit length of 0.35 nm, The *R*_F_ of PEG1000, PEG3000, and PEG5000 are 2.3, 4.4, and 6.0 nm, respectively. Our data show that PEG1000 is already sufficient to lift the bilayers for part of the three proteins to diffuse rapidly in a free-standing-like membrane environment. Moreover, longer PEG polymer resulted in larger mobile fractions of all membrane proteins.

We point out that the density of PEG cushion was kept identical (0.5% of PEGylated lipid in synthetic liposome) when varying the PEG length. As a result, considering the size of PEG polymers (*R*_F_) differs, the free space between the PEG mushroom pillars varies in the three polymer cushioned membranes. The PEG1000 membrane is expected to have the largest free space, and the PEG5000 membrane has the smallest. The average distance between the PEG polymers can be estimated given the molecular composition and the fraction of PEGylated lipid of the membrane. Unfortunately, such information is difficult to obtain because (1) the molecular composition of plasma membrane vesicle is unknown and (2) the exact level of molecular dilution by mixing synthetic liposomes and plasma membrane vesicles during membrane formation is difficult to estimate. Nevertheless, here we calculate the lowest limit (shortest) of the average PEG distance by ignoring the dilution of plasma membrane vesicle: given a POPC molecule has an area of 0.7 nm^2^, the density of PEGylated lipid is one per 11.8 × 11.8 nm^2^ (0.5% of PEGylated lipid). The resulting surface coverage of the PEG (assuming a spherical shape) is 11.9, 43.7, and 81.2% for PEG1000, PEG3000, and PEG5000 membranes. The real density of PEGylated lipid in our plasma membrane bilayers is expected to be lower due to the dilution of plasma membrane vesicles, and thus the real surface coverage should be further reduced. We note that the surface coverage of PEG1000 membrane is quite low (about 10%). It could be that part of the PEG1000 membrane collapses on the solid substrate (uncushioned), resulting in the immobilization of membrane proteins. Further investigation is needed to evaluate the effect of surface coverage of PEG cushion more quantitatively.

### Transient confinements of membrane proteins

To test whether the PEG cushion or any other membrane organization disturbs the diffusion of membrane proteins, we analyze the trajectories of mobile membrane proteins and quantify the probability of apparent transient confinements (Methods). For dye-labeled phospholipid ATTO532-DOPE, the probability of transient confinement, defined as the ratio of total residence time of all detected transient confinements to the overall observation time of all mobile molecules, is low, no more than 5 % in all membrane samples (see Table 1 for the data of confinement probability of ATTO532-DOPE and other membrane proteins). Increasing the PEG length reduces the confinement probability of ATTO532-DOPE, from 4.8 % of PEG1000 membranes to 0.8 % of PEG5000 membranes. Great reduction of confinement probability was also observed in GPI-GFP, CD98-GFP and SSTR3-GFP whose confinement probabilities were 5.1%, 16.2%, and 12.3% in PEG1000 membranes, and they were reduced to 3.3 %, 5.1 %, and 3.0 % in PEG5000 membranes, respectively (see Table 1 for more data). The decrease of transient confinement agrees with and thus may partly explain the enhanced diffusion coefficient measured in membranes with longer PEG lengths. We confirmed that the false detection of transient confinements is rare (around 0.2 %) by applying the same analysis to simulated Brownian motion with the same localization error and trajectory length, indicating the transient confinements detected from membrane proteins are largely real. As an example, Figure 3a plots the diffusion trajectories of GPI-GFP in PEG3000 membrane where transient confinements were highlighted. The size of transient confinements of GPI-GFP in PEG3000 membranes is 104 ± 39 nm (Figure 3b), directly determined from the trajectories. The characteristic residence time τ is 0.07 ± 0.01 s, estimated by an exponential fit of 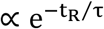 to the histogram (Figure 3c).

**Figure 3.**
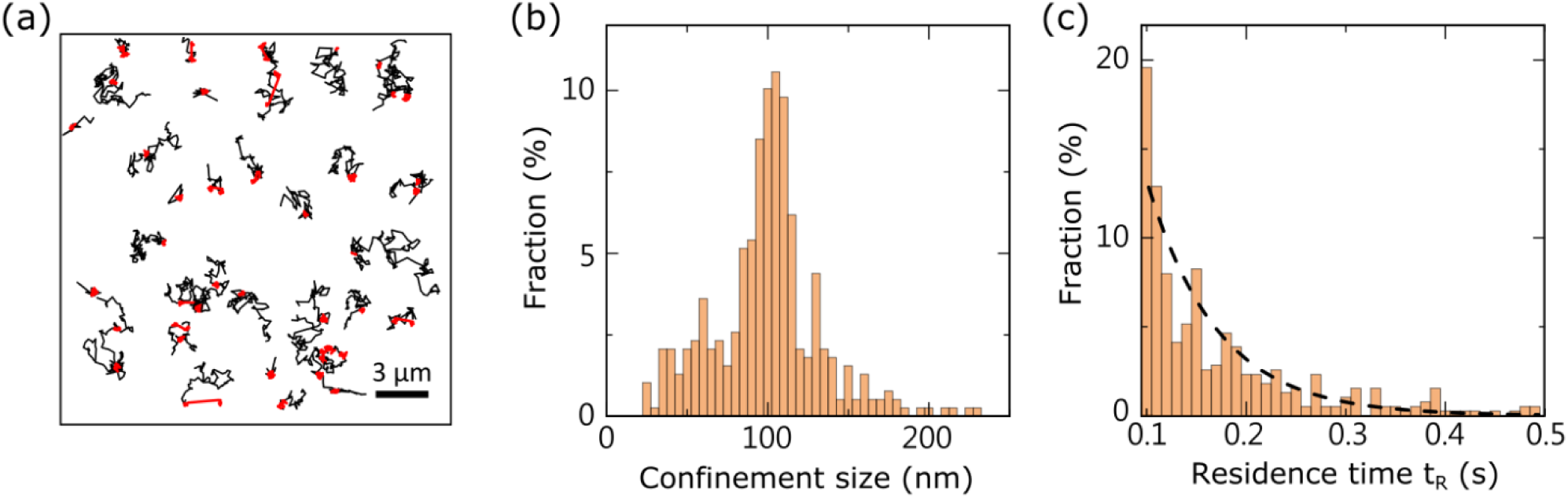
Transient confinements of GPI-GFP in polymer cushioned plasma membrane bilayers (PEG3000 membranes). (a) Diffusion trajectories of single CD98-GFP where transient confinements are highlighted in red. (b) Histogram of transient confinement sizes. (c) Histogram of residence time in transient confinements. The dashed line shows the exponential fit.

### Effect of cholesterol

Membrane molecules may form clusters by self-assembly.^67^ For example, GPI-GFP is well recognized to associate with lipid rafts which are cholesterol enriched membrane nanodomains.^68^ It is possible that the GPI-GFP is attached with cholesterol, sphingolipids, and other raft-associated proteins, and diffuses together as a small complex. The complex may be immobilized once its size becomes too large, and the associated GPI-GFP would appear to be stationary. In addition to GPI-GFP, CD98 was also reported to be associated with lipid rafts.^69^ Herein, we examine the effect of cholesterol by depleting it by chemical reagent (Methods) and quantify the resulting differences in molecular dynamics.

We characterized diffusion of all membrane molecules in our polymer cushioned plasma membrane bilayers in response to cholesterol depletion by treating MβCD for 15 minutes (Methods).^58^ The diffusion coefficient of non-raft phospholipid, ATTO532-DOPE, was almost unchanged by cholesterol depletion (see Table 1 and Figure 4a). We then examined the two raft-associated proteins GPI-GFP and CD98-GFP after cholesterol depletion (Figure 4b and 4c). Surprisingly, the diffusion characteristics of the two proteins remained largely the same after cholesterol depletion, regardless of the polymer lengths. Furthermore, cholesterol depletion did not enhance the mobile fraction of the proteins, either. The fact that cholesterol depletion did not affect measured diffusion characteristics indicates that lipid rafts play a negligible role in the molecular dynamics of the spatial and temporal scale of our measurements, i.e., in the step size of hundreds of nanometers and a time resolution of a few milliseconds. Cholesterol depletion may remove the putative cholesterol-enriched nanodomains that raft-philic proteins were associated to, but such size reduction was hard to be detected from the diffusion in our polymer cushioned membranes (where diffusion coefficient is almost insensitive to the size of membrane inclusion). We also measure the dynamics of SSTR3-GFP before and after cholesterol depletion, and once again, no statistically significant difference was observed (Figure 4d). We further quantified the probability of transient confinements for the four membrane molecules in plasma membrane bilayers after cholesterol depletion. We found that the confinement statistics remained nearly unchanged, indicating that the mechanism of detected transient confinement is independent of cholesterol and thus independent of lipid rafts (see Table 1 for the data).

**Figure 4.**
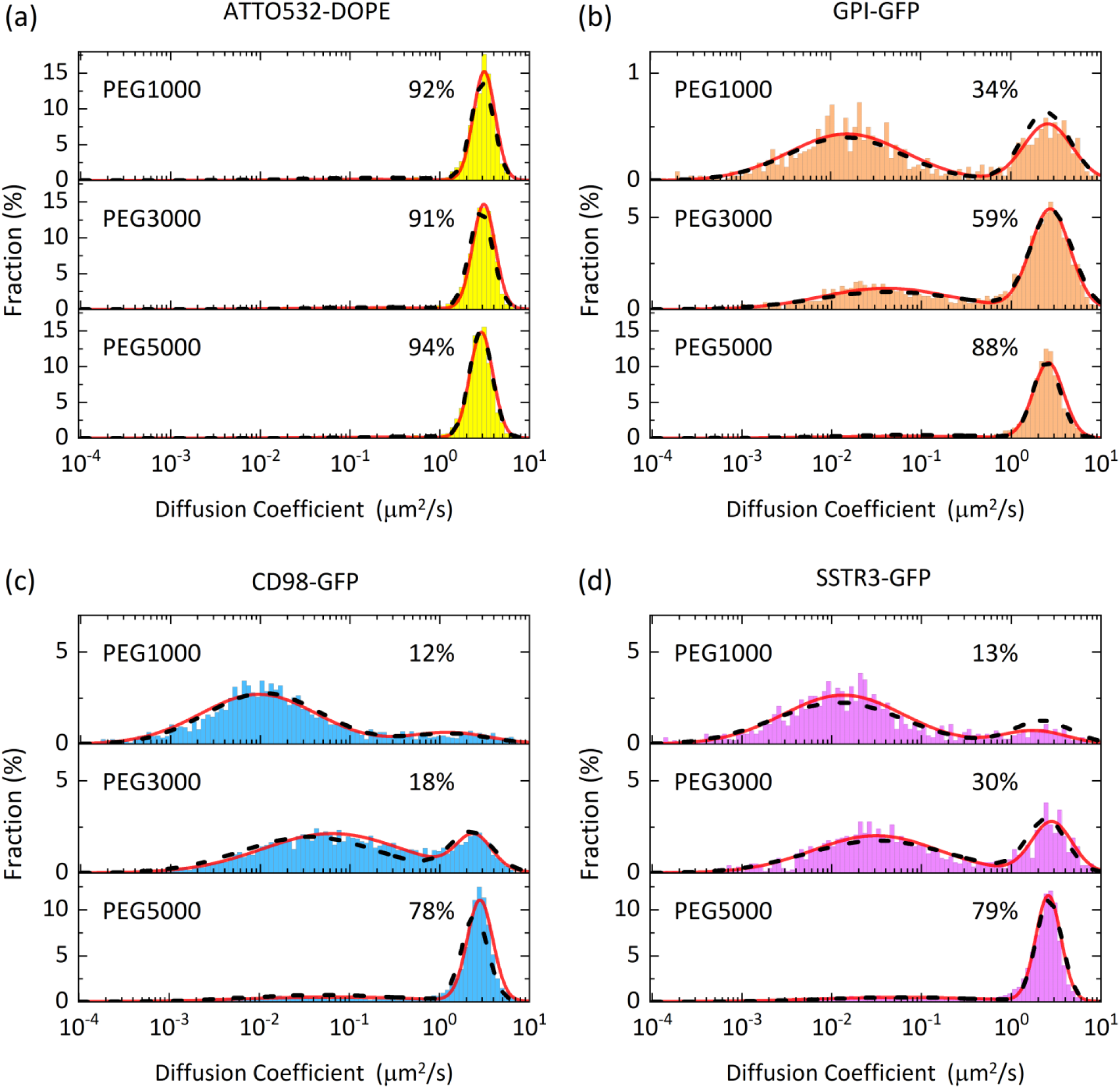
Histograms of diffusion coefficient of four membrane molecules (a) ATTO532-DOPE, (b) GPI-GFP, (c) CD98-GFP, and (d) SSTR3-GFP in polymer cushioned plasma membrane bilayers after cholesterol depletion. Histograms were fitted by a superposition of two Gaussian functions, plotted in red solid lines. For comparison, the results measured before cholesterol depletion (as shown in Figure 2) are plotted in black dashed lines. The percentage displayed in each histogram marks the mobile fraction of the molecule.

## DISCUSSION

Figure 5 summarizes the diffusion characteristics (diffusion coefficient, mobile fraction, and confinement probability) of the four membrane molecules in all membranes prepared in this work. Our data show that the diffusion coefficients of ATTO532-DOPE and GPI-GFP were nearly unchanged by increasing the PEG length, whereas the diffusion coefficients of transmembrane proteins CD98-GFP and SSTR3-GFP were raised by increasing the PEG lengths (Figure 5a). In PEG5000 membranes, all three membrane proteins diffused at a comparable diffusion coefficient of 2.5 μm^2^/s approximately, regardless of their distinct physical sizes. It suggests that the polymer cushion PEG5000 was able to lift the bilayer sufficiently so that the membrane proteins experienced a free-standing-like membrane. With shorter PEG lengths, the diffusion was slowed down most likely due to the interaction with the substrate. The slowdown at short PEG lengths was not observed in ATTO532-DOPE and GPI-GFP possibly because they both resided in the distal membrane leaflet that was readily away from the substrate.

**Figure 5.**
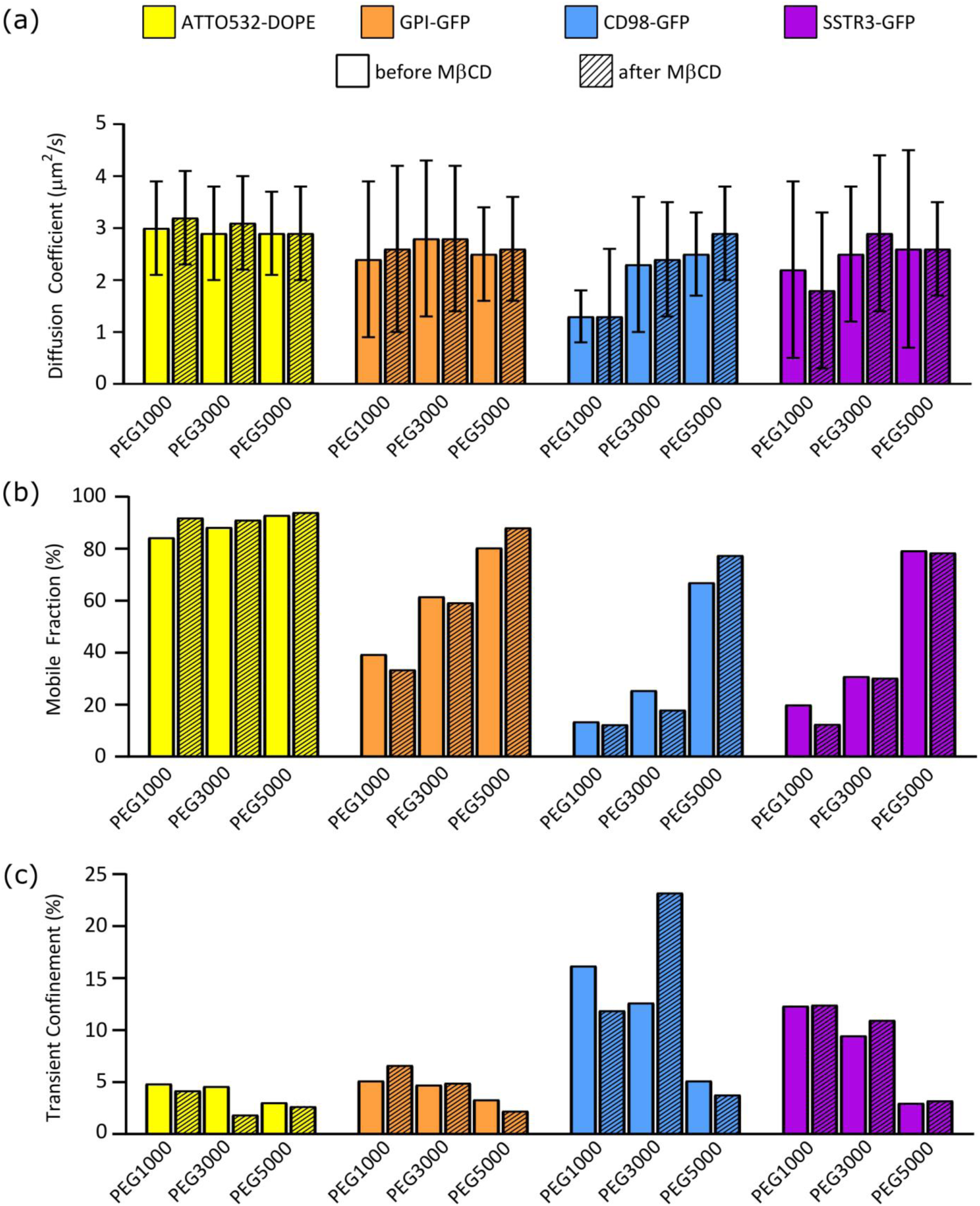
Bar plots of diffusion characteristics of ATTO532-DOPE, GPI-GFP, CD98-GFP, and SSTR3-GFP in polymer cushioned plasma membrane bilayers. (a) Diffusion coefficient of the mobile population. (b) Mobile fraction. (c) Probability of transient confinement.

Increasing the PEG length consistently enhanced the mobile fractions of the three membrane proteins (Figure 5b). Our results agree with previous evidence that increasing PEG lengths helps diminish the interaction of the membrane proteins with the supporting substrate and thus enhances the mobile fractions of membrane proteins.^22, 48^ The two transmembrane proteins (CD98-GFP and SSTR3-GFP) showed a significant enhancement of mobile fraction (> 40%) when the PEG length was increased from PEG3000 to PEG5000. The drastic amplification of the mobile fraction suggests that there is a critical PEG length for creating adequate space between the bilayer and the substrate to accommodate the transmembrane proteins. On the other hand, the mobile fraction of GPI-GFP was improved steadily by increasing PEG length without a sudden enhancement possibly because GPI-GFP is a lipid-anchored peripheral protein whose interaction with the substrate is inherently different from those of transmembrane proteins. Since GPI-GFP does not have cytoplasmic domains, its interaction with the substrate is expected to be indirect—GPI-GFP may appear immobile when being trapped by or associated with other membrane molecules that are pinned on the substrate. Therefore, our observation of the rescue of immobilized GPI-GFP by increasing PEG length mainly reflects the general reduction of membrane-substrate interaction.

Figure 5c summarizes the statistics of transient confinement measured in the four membrane molecules. Increasing the PEG length reduced the probability of transient confinement, indicating the confinement was caused by direct or indirect interaction with the underlying substrate. The two transmembrane proteins (CD98-GFP and SSTR3-GFP) showed much higher tendency of transient confinement than ATTO532-DOPE and GPI-GFP, especially in membranes of shorter PEG lengths (PEG1000 and PEG3000). It implies that the confinements of CD98-GFP and SST3-GFP were due to the interaction between their cytoplasmic domains and the substrate. By increasing the PEG length to PEG5000, the probability of transient confinement became more comparable between the four molecules (3–5%).

We did not see consistent effects of cholesterol depletion to the diffusion characteristics of the four membrane molecules, which is unexpected especially for GPI-GFP and CD98-GFP that were found to be associated with cholesterol-dependent membrane nanodomains (lipid rafts) in live cells.^68-69^ We stress that our data does not imply cholesterol-independent molecular dynamics of the four molecules. Instead, it is likely that the effect of cholesterol (or lipid rafts) was not detectable in our polymer cushioned membranes. Cholesterol depletion is thought to disrupt the formation of nanoclusters raft-associated molecules,^58^ and thus may affect the molecular dynamics. However, it is worth noting that our polymer-cushioned plasma membrane bilayers were free-standing-like membranes (see before-mentioned discussion) where the diffusion coefficient was weakly dependent on the size of membrane inclusion (based on Saffman–Delbrück model).^64^ As a result, although cholesterol depletion could modify the nanoscale membrane organization, the resulting difference in diffusion characteristics may be difficult to detect.

The accuracy of diffusion characterization depends critically on the quality of single-molecule data. Our estimation of diffusion coefficient, mobile fraction, and transient confinement can be more accurate when data of longer trajectories and higher localization precision are available.^60-61^ In this work, the trajectory length and localization precision were primarily limited by the photostability of GFP. It has been demonstrated that diffusion of single membrane molecules can be measured at much higher spatial precision (nanometers) and temporal resolution (microseconds) by using nanoparticles as labels.^26-27^ By using interferometric scattering (iSCAT) microscopy,^70^ very small nanoparticles (as small as 10 nm gold nanoparticle) can be visualized and tracked at a high speed over a long observation time.^71^ The higher spatiotemporal resolution can potentially reveal nanoscopic molecular dynamics in the polymer-cushioned plasma membrane bilayers.

## CONCLUSIONS

We studied dynamics of single membrane proteins in PEG polymer cushioned plasma membrane bilayers by fluorescence microscopy. Four membrane molecules were investigated, including a phospholipid (ATTO532-DOPE), a GPI-anchored protein (GPI-GFP), a single-pass transmembrane protein (CD98-GFP), and a seven-pass transmembrane protein (SSTR3-GFP). Their dynamics were measured in three different membrane systems with various PEG lengths (PEG1000, PEG3000, and PEG5000). Large amount of data were collected for each molecule in every membrane conditions in an automated manner. We found that introduction of PEG with increasing length (from PEG1000 to PEG5000) drastically enhanced the mobile fraction of membrane molecules. The diffusion coefficients of transmembrane proteins (CD98-GFP and SSTR3-GFP) were also increased by polymer cushion with increasing PEG length. However, diffusion coefficients of lipid and lipid-anchored protein (ATTO532-DOPE and GPI-GFP) remained unchanged in membranes of different cushions. The polymer cushion of increasing PEG length also reduced the probability of transient confinements of membrane molecules. By modulating the concentration of cholesterol in the plasma membrane bilayers, we concluded that cholesterol plays a negligible role in all measured dynamics, and thus no effect of lipid rafts was detected in our experiments. Our characterization of membrane protein dynamics elucidated the biophysical properties of polymer cushioned plasma membrane bilayers that are valuable for future applications in membrane-based biosensors and analytical assays.

## Supporting Information

Single-molecule fluorescence videos of GPI-GFP (Movie S1), CD98-GFP (Movie S2), SSTR3-GFP (Movie S3), and ATTO532-DOPE (Movie S4) in PEG1000 membranes; fluorescence confocal microscope images of cells transfected by the plasmids of GFP-fused membrane proteins (Figure S1); size distribution of cell blebbed vesicles and synthetic liposomes (Figure S2).

## ACKNOWLEDGMENTS

This project was financed by the Career Development Award of Academia Sinica (AS-CDA-107-M06) and Ministry of Science and Technology, Taiwan (MOST 105-2112-M-001-016-MY3). We thank Hsiao-Chieh Chen for technical support.

